# Updated Biomarkers for TNBC in African vs. Caucasian American Women

**DOI:** 10.1101/2023.07.17.549415

**Authors:** Jacob Croft, Odalys Quintanar, Jun Zhang

## Abstract

**Introduction:** Breast cancer, especially triple-negative breast cancer (TNBC), is a significant concern in the US, being the most common cancer among women and the second leading cause of cancer-related deaths. TNBC lacks crucial receptors targeted in other breast cancer types, leading to a poor prognosis and limited treatment options due to its aggressive and heterogeneous nature.

**Significance:** African American women (AAW) with TNBC face higher mortality rates and more aggressive disease compared to Caucasian American women (CAW). Despite efforts to find biomarkers specific to AAW and CAW with TNBC, limited sample availability and data resources have been obstacles.

**Methods:** In our study, we examined 237 candidate peptide biomarkers using publicly available data.

**Results:** identify 23 unique prognostic biomarkers. These biomarkers accurately assess patient conditions based on race-specific gene expression patterns, holding potential to address racial disparities in TNBC treatment.

**Conclusion:** Overall, our research sheds light on how racial genetic profiles influence TNBC prognosis and treatment efficacy. The identified prognostic biomarkers pave the way for future studies addressing TNBC racial disparities and personalized treatment approaches.

## Introduction

Breast cancer is the primary type of cancer affecting women in the United States, representing 30% of newly diagnosed cases and ranking as the second leading cause of cancer-related death among women [1-5]. The prognosis of breast cancer depends heavily on factors such as grade, subtype, and early detection. Among the various subtypes, triple-negative breast cancer (TNBC) has the poorest overall prognosis [6-8]. TNBC is an aggressive subtype characterized by the absence of three receptors (ER, nPR, HER2) [6,9,10]. It displays heterogeneity, leading to uncertain outcomes [5,11-13]. Treatment options for TNBC are limited, and its aggressive nature is accompanied by a survival disparity that is not fully understood between Caucasian and African American women [3,5,9,12,14-16]. African American women (AAW) face disparities in breast cancer, experiencing more aggressive forms of the disease and higher mortality rates, particularly in cases of TNBC [9,17-22].

The incidence of TNBC is higher in African American women (AAW) compared to Caucasian American women (CAW) [23]. AAW diagnosed with breast cancer face nearly 40% higher mortality rates, which can be influenced by socioeconomic factors and genetic predispositions [23]. Studies have shown that AAW with breast cancer have approximately 40% higher mortality rates than CAW of similar age and prognosis [18,24-29]. Even when receiving equal treatment, AAW with TNBC experience a worse prognosis [24-26,28,30]. Various factors such as differences in mutational landscapes, gene expression, and tumor microenvironment contribute to the poor outcomes observed in AAW with TNBC [25,30-36]. The higher prevalence of TNBC and lower survival rates in young AAW suggest the presence of race-associated molecular mechanisms in breast cancer [37-40].

The diverse clinical characteristics and treatment responses observed in breast cancers, particularly TNBCs, underscore the importance of identifying specific biomarkers for each subtype to inform clinical decision-making [41-46]. Breast cancers, especially TNBCs, exhibit diverse clinical characteristics and treatment responses, highlighting the need for identifying specific biomarkers for each subtype. This information becomes crucial for guiding clinical decisions due to the heterogeneity of TNBCs [41-46]. Despite this challenge, the heterogeneity also presents an excellent opportunity to discover potential diagnostic and prognostic biomarkers. Biomarkers play a crucial role in providing valuable insights into cancer staging, location, and signaling pathways [11,47-54]. Numerous studies have emphasized the significance of biomarkers in comprehending the various TNBC subtypes, highlighting both the advantages and disadvantages of Lehmann’s classification system [5,53,55]. With advancements in technology and expression profiling, our understanding of TNBC subgroups has been significantly enhanced [56-59], allowing for the identification of precise biomarkers, therapeutic targets, and molecular mechanisms associated with the disease [60-63].

Consequently, there exists an unmet need to create a comprehensive collection of potential prognostic biomarkers that are specifically associated with AAW-TNBCs and CAW-TNBCs. These biomarkers would not only serve as a reliable prognostic tool but also advance our understanding of their unique molecular profiles and clinical implications. Numerous research groups, including our own, have dedicated significant efforts towards identifying a panel of prognostic biomarkers specifically associated with AAW-TNBCs and CAW-TNBCs, yielding some noteworthy progress. However, these studies face evident limitations primarily due to the extremely small number of AAW-TNBC cases available for prognostic marker determination, as well as the limited availability of publicly accessible gene expression profiles and datasets that predominantly consist of tumors from Caucasian individuals. Therefore, the utilization of this data to address racial disparities has consistently faced scrutiny and encountered challenges, as highlighted by the arguments mentioned earlier. These issues have significantly impeded the progress of identifying and validating candidate biomarkers that possess specific clinical utility for African American patients.

## Materials and Methods

The TNBC patient data for this study were obtained from a major TNBC public repository, the TCGA (The Cancer Genome Atlas) database for Breast Invasive Cancer (BRCA) patients, with recent major inputs from The University of Alabama at Birmingham Cancer data portal (UALCAN) [64,65]. The study included a total of 1041 subjects, categorized as normal (n=114), CAW (Caucasian American women, CAW) (n=748), and AAW (African American women, AAW) (n=179). Analysis was conducted based on a list of known biomarkers established by our lab in previous studies [66-72], including our own contributions [73-75].

Next, we examined each candidate biomarker by comparing their TPM (transcripts per million) values among three racial groups (African American, Caucasian American, and Asian/Pacific) and a control from the database. This comparison allowed us to identify a more reliable set of biomarkers. Statistical significance between the expression values of the groups was determined using a student’s T test. We specifically looked for differences in expression between the control group, AAW, and CAW. Finally, the selected candidate biomarkers with significantly differential TPM expression levels between the groups were further analyzed in relation to the Kaplan-Meier (KM) survival curves. To create individual KM curves, we utilized the log-rank test to establish unbiased nonparametric categories before comparing them. By using this approach, we not only identified group expression differences but also understood their impact on patients’ prognosis. This allowed us to discover unique prognostic biomarkers that could serve as race-specific TNBC biomarkers [73,74].

## Results

### Over 200 AAW- and CAW-INBC candidate biomarkers identified

Over the past decade, significant efforts have been made to identify prognostic biomarkers specific to AAW-TNBCs and CAW-TNBCs, resulting in the discovery of over 200 potential peptide biomarkers [67-75], and even encompassing metabolites and non-coding RNAs [76,77]. Despite this progress, these previous studies have faced considerable limitations, primarily due to the scarcity of AAW-TNBC cases, which hinders the determination of prognostic markers. To comprehensively address this concern, we capitalized on the rapid growth of published research and publicly available data.

We conducted a collective examination of 237 candidate peptide biomarkers previously reported to evaluate their prognostic potentials. This evaluation was based on combined criteria, including differential expression levels and their impact on patients’ survival rates.

### Validation of 23 AAW-specific prognostic biomarkers

To ensure the accuracy of these biomarkers, we followed a rigorous two-step test process. Firstly, we confirmed that one of the two racial groups showed statistically significant differences from the normal values. Once this was established, we further analyzed the TPM values to find statistical differences between the two racial groups (Figure 1A). Next, we proceeded to examine the expression changes to assess their influence on group prognosis. The KM survival models revealed significant statistical differences between the survival curves of AAW and CAW (Figure 1B). Through this comprehensive two-step examination, we successfully identified 23 distinct and currently validated prognostic biomarkers. These biomarkers offer precise assessments of patient-specific conditions, as they are influenced by unique gene expression patterns specific to each racial group. Although some of the 237 candidate peptide biomarkers showed differences in KM survival outcomes, we couldn’t exclude the possibility of social-economic factors influencing these results without considering TPM expression differences. In conclusion, our comprehensive evaluation of both differential expression levels and their impact on patients’ survival rates led to the validation of twenty-three biomarkers from the current repertoire database (Table 1).

**Table 1.** Summary of selected biomarkers with statistically significant difference to distinguish CAW- and AAW-TNBC Patients. Through our analysis, we have identified 23 distinct AAW- and CAW-TNBC prognostic biomarkers that exhibit significant effectiveness in a large-scale model. The biomarkers highlighted in red are for highly expression levels, while colored in blue for low/medium expression levels. Similarly, the red color-coded gene symbol of biomarkers indicate that the expression patterns are associated with a negative impact on AAW-TNBC patient prognosis, while blue colored genes suggest with a negative impact on CAW-TNBC patient prognosis.

| Gene Name | Race | TPM | AAW v CAW P-value | Ctrl v AAW | Ctrl v CAW | Survival P-value |
| --- | --- | --- | --- | --- | --- | --- |
| AURKB | AAW | 21.527 | 2.6147E-11 | 1.6244E-12 | 1.1102E-16 | 0.034 |
|  | CAW | 9.514 |  |  |  |  |
| PAQR6 | AAW | 5.025 | 2.0619E-06 | 1.6253E-12 | <1E-12 | 0.029 |
|  | CAW | 2.76 |  |  |  |  |
| CDC20 | AAW | 39.097 | 4.257E-08 | <1E-12 | 1.6245E-12 | 0.0094 |
|  | CAW | 22.287 |  |  |  |  |
| HPR | AAW | 0 | 3.6964E-02 | 2.6132E-01 | 6.4190E-04 | 0.034 |
|  | CAW | 0 |  |  |  |  |
| GPT2 | AAW | 15.668 | 3.7964E-03 | 2.2780E-02 | 1.2479E-06 | 0.041 |
|  | CAW | 12.066 |  |  |  |  |
| ATP6V1C2 | AAW | 2.076 | 1.1280E-02 | 1.4163E-06 | 1.5838E-04 | 0.0066 |
|  | CAW | 1.47 |  |  |  |  |
| ITPA | AAW | 76.386 | 6.7734E-12 | <1E-12 | 1.6245E-12 | 0.042 |
|  | CAW | 60.515 |  |  |  |  |
| UQCRB | AAW | 58.925 | 1.8224E-03 | 8.0222E-05 | 1.0626E-01 | 0.00023 |
|  | CAW | 52.536 |  |  |  |  |
| CHCHD1 | AAW | 81.589 | 1.7681E-07 | 1.6244E-12 | <1E-12 | 0.039 |
|  | CAW | 70.837 |  |  |  |  |
| RPP21 | AAW | 95.852 | 1.6284E-12 | 1.6245E-12 | 1.6245E-12 | 0.00023 |
|  | CAW | 65.736 |  |  |  |  |
| POMC | AAW | 0.766 | 3.7768E-03 | 2.5028E-03 | 9.6602E-01 | 0.0053 |
|  | CAW | 0.341 |  |  |  |  |
| LINGO3 | AAW | 0.143 | 5.2287E-04 | 1.0938E-09 | 6.7057E-14 | 0.0021 |
|  | CAW | 0.095 |  |  |  |  |
| AGAP4 | AAW | 7.092 | 1.0828E-03 | 4.1088E-01 | 1.1360E-02 | 0.05 |
|  | CAW | 5.69 |  |  |  |  |
| BCL2L12 | AAW | 17.806 | 1.5909E-13 | <1E-12 | <1E-12 | 0.016 |
|  | CAW | 12.436 |  |  |  |  |
| OBSL1 | AAW | 57.571 | 1.1904E-04 | 4.3049E-03 | 2.1560E-01 | 0.036 |
|  | CAW | 44.276 |  |  |  |  |
| ZNRD1 | AAW | 53.11 | 4.3514E-03 | 1.6246E-12 | 1.6246E-12 | 0.002 |
|  | CAW | 44.415 |  |  |  |  |
| MRPL18 | AAW | 89.465 | 6.0074E-04 | 1.6244E-12 | <1E-12 | 0.00031 |
|  | CAW | 80.863 |  |  |  |  |
| PDCD10 | AAW | 54.417 | 3.965E-04 | 3.0824E-08 | 1.6244E-12 | 0.046 |
|  | CAW | 62.289 |  |  |  |  |
| DDX1 | AAW | 71.496 | 1.834E-07 | 1.5090E-08 | 4.0598E-01 | 0.032 |
|  | CAW | 79.725 |  |  |  |  |
| LDHA | AAW | 399.349 | 7.8349E-07 | 2.9809E-01 | <1E-12 | 0.024 |
|  | CAW | 488.868 |  |  |  |  |
| SEPT7 | AAW | 101.467 | 1.6247E-12 | 1.6244E-12 | 1.0403E-04 | 0.025 |
|  | CAW | 128.783 |  |  |  |  |
| IKBIP | AAW | 14.176 | 8.2435E-04 | 1.9356E-05 | 3.5997E-02 | 0.037 |
|  | CAW | 16.732 |  |  |  |  |
| SIK1 | AAW | 19.072 | 1.892E-01 | 4.1577E-02 | 1.7401E-05 | 0.02 |
|  | CAW | 17.116 |  |  |  |  |

**Figure 1:**
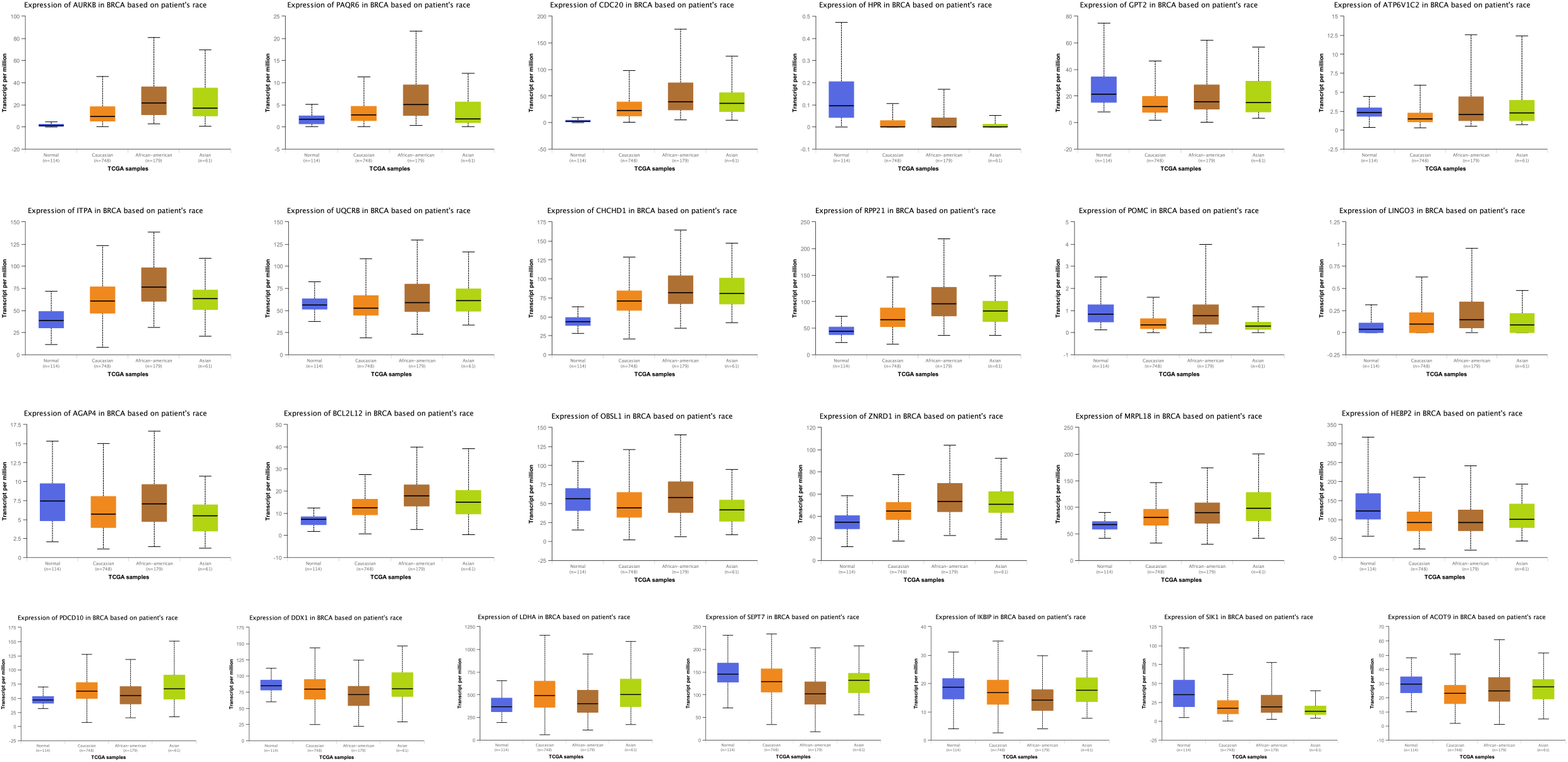
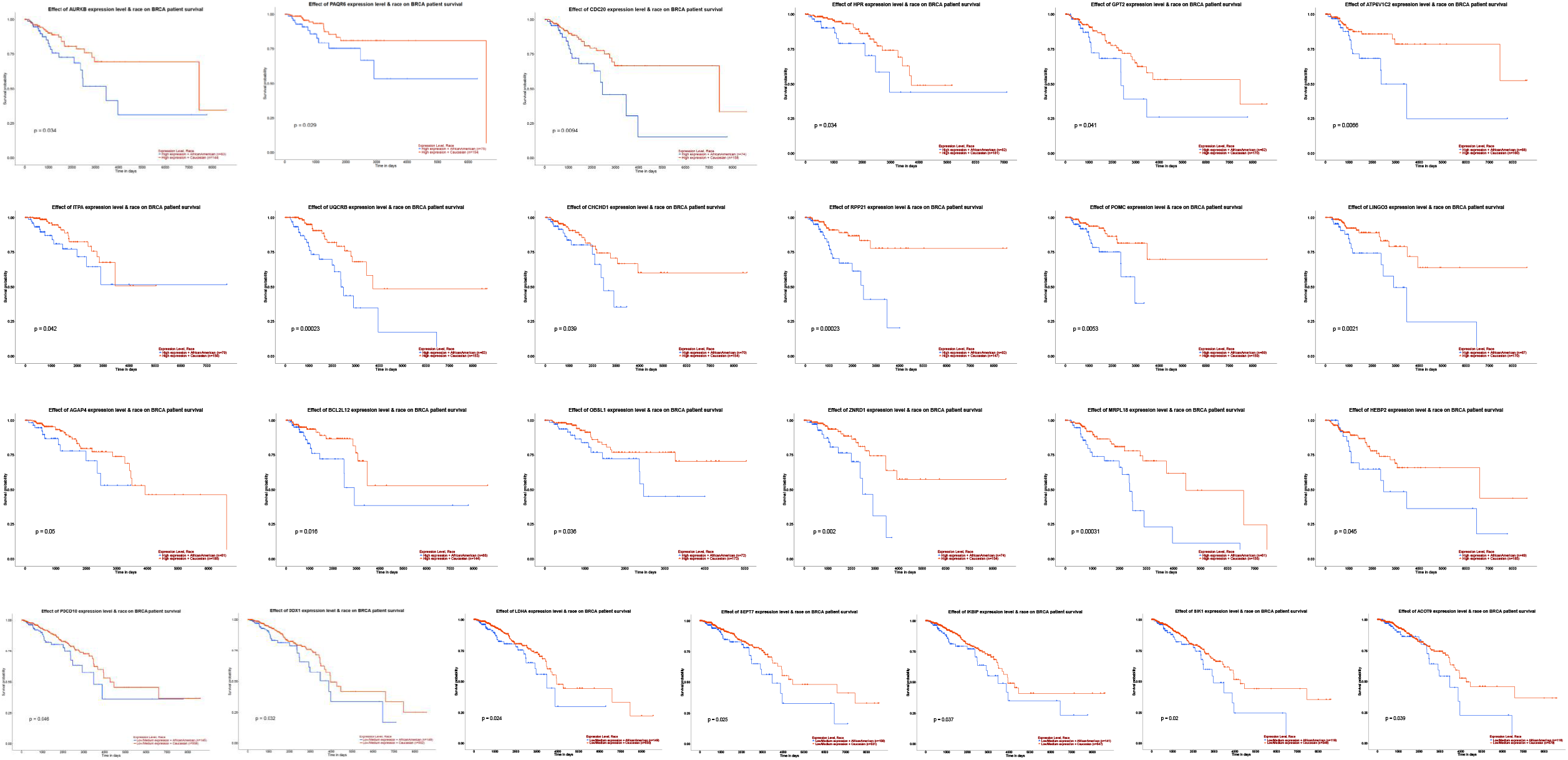
Statistical significant different expression and survival curves for novel and candidate biomarkers based on racial cancer disparity between AAW- and CAW-TNBCS. Prognostic effects of genes based on expression level and patients’ rate on the survival rate. Using the UALCAN portal, we accessed the TCGA database for BRCA (Breast Invasive Cancer) patients and input different genes to be scanned through the database. **1A:** The prognostic impact of biomarkers were assessed based on combined evaluation of their differential expression levels and how they affected the survival rate of patients. To conduct the analysis, the UALCAN portal provided access to the TCGA database containing data on TNBC patients. Previously reported biomarkers were selected for examination, and their expressions were queried within the database. Genes were measured in TPM and compared using a student T test to establish racial cancer disparities. **1B:** Prognostic effects of genes based on expression level and patients’ rate on the survival rate. Using the UALCAN portal, we accessed the TCGA database for BRCA (Breast Invasive Cancer) patients and input different genes to be scanned through the database. The UALCAN generated the KM survival curves for each gene. The curves in the top row are all for genes that are shown to have a significant difference in survival probability between AAW and CAW when in high expression. The bottom two rows are comprised of curves for genes that had a significant difference in AAW and CAW survival rates when in low/medium expression.

## Discussion

The study aimed to investigate how racial genetic profiles can impact prognosis and treatment effectiveness, taking into account the influence of socioeconomic status (SES) on treatment disparities. Current biomarkers were used to gain a deeper understanding of these factors in patient outcomes. We established two criteria for evaluating the efficacy of race-specific biomarkers. First, we compared TPM values to identify expression differences between the control and racial groups, as well as between the two racial groups. Then, we analyzed the survival curves of these genes to assess differences in prognostic outcomes. This two-tiered approach ensured a robust establishment of biomarker status while considering SES-related factors. The study’s successful foundation has provided valuable insights into these complex relationships.

The study also highlighted the significance of racial differences in prognosis and occurrence rates, underscoring the need to discover more needed prognostic biomarkers. Through the assessment of 237 candidate peptide biomarkers, we successfully identified 23 prognostic biomarkers, laying the groundwork for a potential prognostic tool to address TNBC racial disparities. Additionally, this research uncovered potential genetic mechanisms contributing to the development of the disease, offering new avenues for TNBC treatment options.

Breast cancer treatment often involves hormone therapy due to the influence of steroid hormones. However, TNBC, lacking estrogen and progesterone receptors, presents a challenge for conventional treatment. Nevertheless, the discovery of the CmP signal networks associated with progesterone signaling offers new opportunities for TNBC therapies [73,74,78,79]. In this study, we validated certain biomarkers, such as PDCD10 and PAQR6, key components of the CmP signal network, as promising prognostic indicators for AAW- and CAW-TNBC. These findings support our argument and align with our previous research. Additionally, they may have implications for immune-subtyping, similar to observations in liver cancers. However, more research is needed to fully explore their potential and develop targeted treatments for TNBC.

## Conclusion

Our research provides valuable insights into the influence of racial genetic profiles on TNBC prognosis and treatment efficacy. The identified prognostic biomarkers offer a solid foundation for future studies addressing TNBC racial disparities and may lead to personalized treatment approaches.

## Acknowledgments

not applicable.

## Statement of Ethics

not applicable.

## Author Contributions

Jacob Croft: Writing-Original draft preparation, Writing-Reviewing and Editing, Writing-Reviewing and Editing; Odalys Quintanar: Writing-Original draft preparation, Writing-Reviewing and Editing; Jun Zhang: Conceptualization, Methodology, Writing-Original draft preparation, Writing-Reviewing and Editing.

## Data Availability Statement

not applicable.

