## Supplementary material for "Updated Biomarkers for TNBC in African vs. Caucasian American Women": Suppl Materials

Running Title: updated TNBC biomarkers between CAW and AAW

**Supplemental Materials**

***Correspondence:***

Jun Zhang, Sc.D., Ph.D.,

Department of Biomedical Sciences

Texas Tech University Health Science Center

5001 El Paso Drive, El Paso, TX 79905

Supplemental Legends:

**Supplemental Figure 1: An assessment of previously established biomarkers to examine gene disparities specific to African American women (AAW) and Caucasian American women (CAW) racial groups.** As a preliminary step in our research, we conducted a re-evaluation of the biomarkers previously established to assess their continued efficacy on a larger scale. (**A**). The first step involved examining the TPM values and making comparisons between different racial groups and a control group unaffected by cancer. Surprisingly, in the larger study, we observed less significant differences in TPM gene expression compared to what was initially found in smaller study populations. (**B**). We then proceeded to analyze the prognostic effect of these same genes using survival curves from the initial studies. Despite some variations in the survival curves, we are no longer inclined to consider these genes as race-specific biomarkers. The reason for this decision is that there were no significant differences between African American women (AAW) and Caucasian American women (CAW) patients, making it challenging to attribute the variations solely to racial factors. Instead, socioeconomic differences might play a more significant role as the causative agent.

### Figure 1A

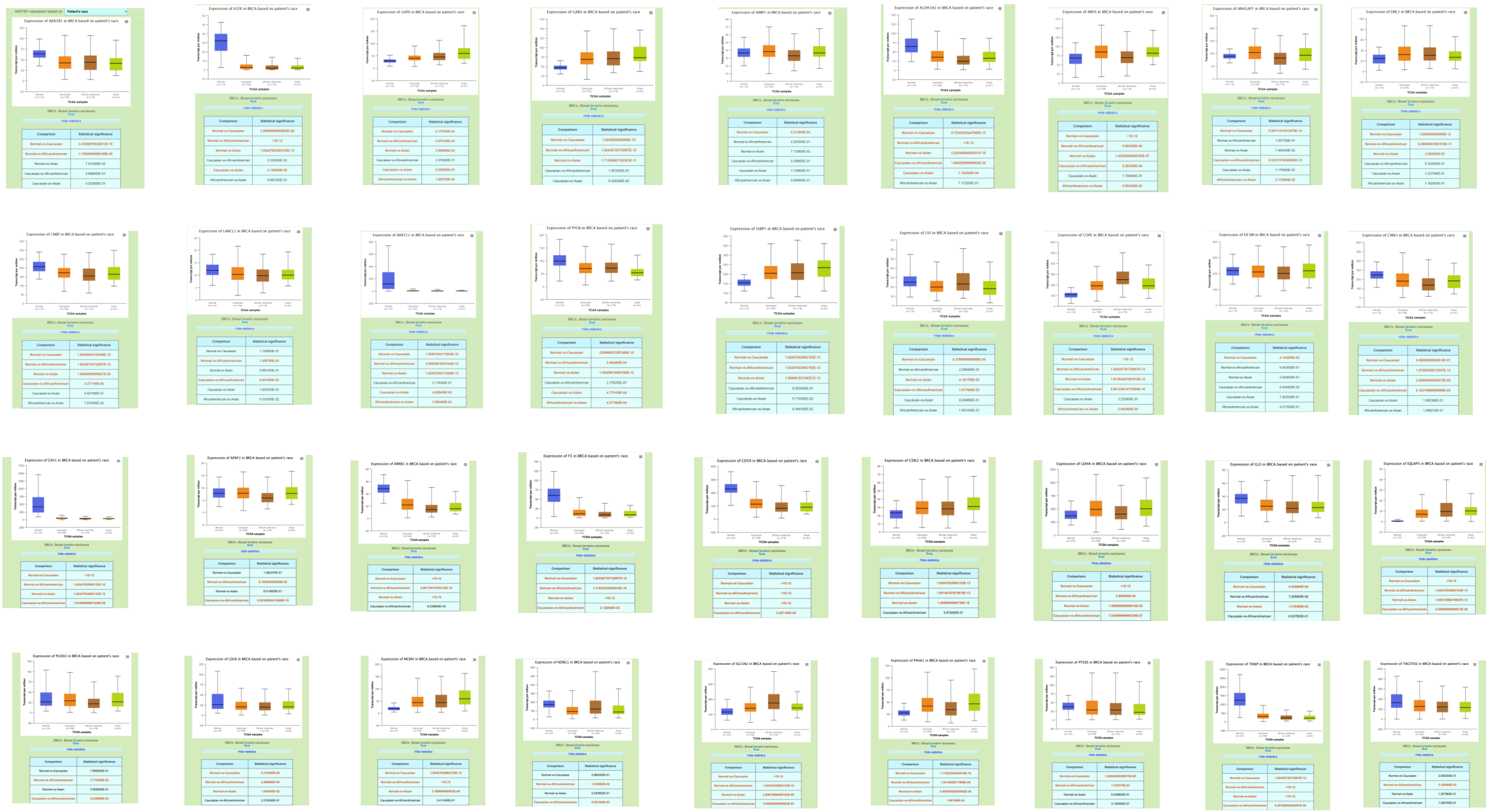

### Figure 1B

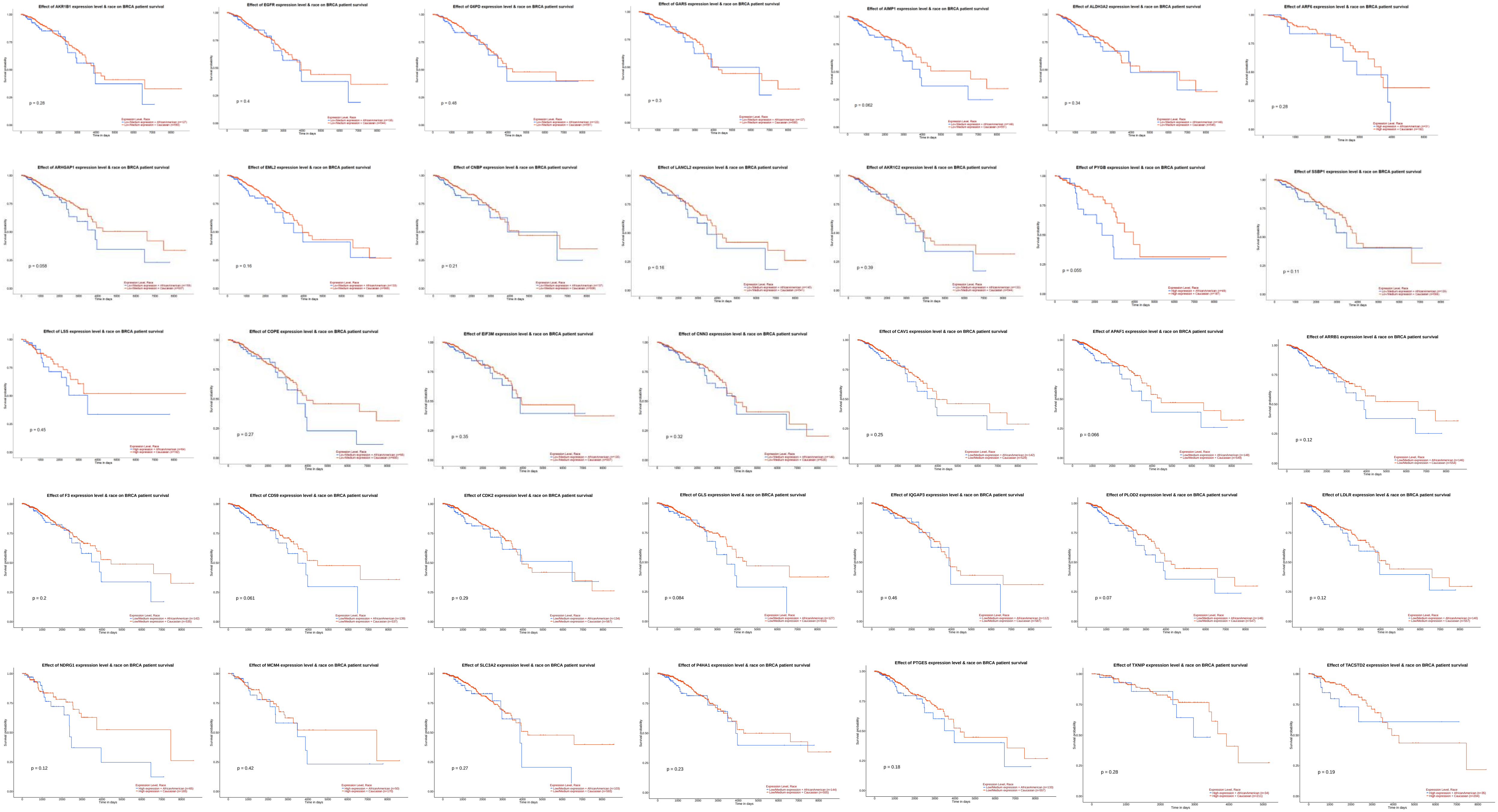
Effect of AKR1B1 expression level & race on BRCA patient survival
p = 0.28
Effect of EGFR expression level & race on BRCA patient survival
p = 0.4
Effect of G6PD expression level & race on BRCA patient survival
p = 0.48
Effect of GARS expression level & race on BRCA patient survival
p = 0.3
Effect of AIMP1 expression level & race on BRCA patient survival
p = 0.062
Effect of ALDH3A2 expression level & race on BRCA patient survival
p = 0.34
Effect of ARF6 expression level & race on BRCA patient survival
p = 0.28
Effect of ARHGAP1 expression level & race on BRCA patient survival
p = 0.058
Effect of EML2 expression level & race on BRCA patient survival
p = 0.16
Effect of CNBP expression level & race on BRCA patient survival
p = 0.21
Effect of LANCL2 expression level & race on BRCA patient survival
p = 0.16
Effect of AKR1C2 expression level & race on BRCA patient survival
p = 0.39
Effect of PYGB expression level & race on BRCA patient survival
p = 0.055
Effect of SSBP1 expression level & race on BRCA patient survival
p = 0.11
Effect of LSS expression level & race on BRCA patient survival
p = 0.45
Effect of COPE expression level & race on BRCA patient survival
p = 0.27
Effect of EIF3M expression level & race on BRCA patient survival
p = 0.35
Effect of CNN3 expression level & race on BRCA patient survival
p = 0.32
Effect of CAV1 expression level & race on BRCA patient survival
p = 0.25
Effect of APAF1 expression level & race on BRCA patient survival
p = 0.066
Effect of ARRB1 expression level & race on BRCA patient survival
p = 0.12
Effect of F3 expression level & race on BRCA patient survival
p = 0.2
Effect of CD59 expression level & race on BRCA patient survival
p = 0.061
Effect of CDK2 expression level & race on BRCA patient survival
p = 0.29
Effect of GLS expression level & race on BRCA patient survival
p = 0.084
Effect of IQGAP3 expression level & race on BRCA patient survival
p = 0.46
Effect of PLOD2 expression level & race on BRCA patient survival
p = 0.07
Effect of LDLR expression level & race on BRCA patient survival
p = 0.12
Effect of NDRG1 expression level & race on BRCA patient survival
p = 0.12
Effect of MCM4 expression level & race on BRCA patient survival
p = 0.42
Effect of SLC3A2 expression level & race on BRCA patient survival
p = 0.27
Effect of P4HA1 expression level & race on BRCA patient survival
p = 0.23
Effect of PTGES expression level & race on BRCA patient survival
p = 0.18
Effect of TXNIP expression level & race on BRCA patient survival
p = 0.28
Effect of TACSTD2 expression level & race on BRCA patient survival
p = 0.19
